# Mapping of Quantitative Trait Locus (QTLs) that contribute to Drought Tolerance in a Recombinant Inbred Line Population of horsegram (*Macrotyloma uniflorum*)

**DOI:** 10.1101/2021.03.09.434553

**Authors:** Megha Katoch, R.K. Chahota

## Abstract

Horsegram (*Macrotyloma uniflorum*) is a drought hardy legume which can be grown in varied soil and temperature regime. It is an important food legume with environmental, nutritive and medicinal benefits. But in terms of genetic improvement it still lags behind other legumes. To get insight into the genetics of tolerance to drought stress, quantitative trait loci for drought tolerance traits were identified using an intraspecific mapping population comprising of 162 F_8_ Recombinant Inbred Lines derived from a cross between HPKM249 and HPK4. A total of 2011 markers were screened on parental lines for polymorphism survey, out of which 493 markers were found to be polymorphic and used for genotyping of the RIL population. Of these 493 polymorphic markers, 295 were assigned to ten linkage groups at LOD 3.5 spanning 1541.7cM with a mean distance of 5.20 cM between adjacent markers. This linkage map along with the phenotypic data for drought tolerance traits was used to identify regions of the horsegram genome in which the genes for the qualitative traits linked to drought tolerance located. A total of seven QTLs were identified for six different drought related traits. One QTL for malondialdehyde content on linkage group 2, two QTLs for root length on linkage group 3 & 9, one QTL each for proline content and chlorophyll content under drought stress on linkage group 4, one QTL each for root dry weight and root fresh weight on linkage group 5 were identified using composite interval mapping. The linkage map and identified QTLs will be utilized in Marker Assisted Breeding and increase our understanding on the physiology of drought stress tolerance. It will also aid in molecular breeding efforts for further genetic improvement of horsegram.

## Introduction

Horsegram (*Macrotyloma uniflorum*) is an useful pulse from prehistoric times which is mainly cultivated for food, feed and medicine by pastoral society in African and Asian continents. It provide an inexpensive source of protein for the people who depends on vegetarian diet. It is classified as Macrotyloma genus, Phaseoleae tribe, Faboideae sub-family, Fabaceae family (Ranasinghe and Ediriweera 2017) and have diploid chromosome numbers 2n = 20 (Cook et al. 2005).The probable genome size is approximately 400 Mbps (Hirakawa et al. 2017). The name *Macrotyloma* is originated from three Greek words *macros, tylos* and *loma* which mean large, knob and margin respectively, representing pod’s knobby statures of the crop (Blumenthal and Staples 1993). The genus Macrotyloma comprises of 32 wild species which are distributed in African, Australian and Indian subcontinent and *Macrotyloma uniflorum* var. *uniflorum* grown in Indian subcontinent is regarded as the only cultivated species. *Macrotyloma uniflorum* var. *uniflorum* is indigenous to parts of Africa and Southeast Asia but Southern India is considered as its centre of origin (Vavilov 1951; Zohary 1970). In India it is cultivated in large area compared to other countries for the purpose of human food and medicine in Ayurveda with the maximum area of cultivation in Andhra Pardesh, Karnataka and Tamil Nadu. It is also grown in many states of India viz. Odisha, West Bengal, Chhattisgarh, Madhya Pradesh, Bihar, Jharkhand, Uttaranchal and Himachal Pradesh. In India cultivation area is 3.126 Lakh ha, production is 1.343 Lakh tonnes with yield of 430 Kg/ha (Directorate of Economics and Statistics (DES), 2016–2017).

Horsegram is a very adaptable crop which can grow from near sea level to 1800m above mean sea level. It can resist drought and wide range of temperature regimes and withstand stress conditions in those areas where other crops are unable to survive (Krishna 2010; Ramya et al. 2013). Being a leguminous crop, it also condition soil with nitrogen and aids in improving the fertility of soil. Along with its comprehensive growing parameters, it has many wide uses such as it can used as green manure, its husks is used for its water holding capacity (Nezamuddin 1970; Zaman and Mallick 1991) and its small height enables it to be sown under taller plant as understory crop (Nezamuddin 1970). It has a high nutritive value and is abundant in protein (17.9 – 25.3%), essential amino acids, carbohydrates (51.9 – 60.9%), iron molybdenum, phosphorus, fibre and various vitamins like Vitamin A, Vitamin B1, Vitamin B2, Vitamin B3 and vitamin C (Sodani et al. 2004). From ancient times it is being used in Ayurveda as an important medicinal crop. Different parts of horsegram plants are used in various trearments viz. seeds of horsegram are used for remedy in urinary stone and piles (Yadava and Vyas 1994), it also act as tonic (Brink 2006), treat problems in menstrual cycle (Neelam 2007) and also used to cure hiccups and worms (Chunekar and Pandey 1998). Also, the gravy of cooked horsegram seeds produces heat that when consumed is used to cure fever, common cold and throat infection (Perumal and Sellamuthu 2007). All the valuable attributes of this legume ensure its cultivation since prehistoric times and make it a promising food source for the future (National Academy of Sciences 1978).

Drought tolerance is a highly complex phenotype. It is affected by many traits such as proficient water uptake through deep roots, reduced rate of transpiration via stomata and limited effect of decreased water potential on metabolites production and plant growth (Manavalan et al. 2009). Plants adapt themselves in drought stress by different mechanisms like drought escape or drought tolerance. Agrarain traits controlling the drought stress tolerance are polygenic in nature and are quantitatively inherited (Gupta et al. 2017). Therefore mapping the quantitative trait locus (QTL) which are linked to drought tolerance-related traits by providing drought condition will aids in the identification of drought-responsive QTLs or genes which can further assist in molecular breeding for traits selection and introgression in breeding strategies through marker assisted selection (MAS). After the discovery of DNA markers and their utilization in the development of framework genetic linkage maps, many QTLs linked to important agronomic traits have been identified (Kim et al. 2015). Although studies on identification of QTLs for drought tolerance in many leguminous crops have been done viz. chickpea (Varshney et al. 2014; Jaganathan et al. 2015), cowpea (Muchero et al. 2009), common bean (Asfaw and Blair 2012; Blair et al. 2012), lentil (Singh et al. 2016) and mungbean (Liu et al. 2017) but not much work has been done in horsegram.

Therefore, study was carried out to construct genetic linkage map of horsegram and to identify the genomic position, number and magnitude of QTLs affecting genetic variation for drought tolerance in RIL populations derived from a cross between two horsegram genotypes (HPKM249 and HPK4) showing contrasting expression for some drought-related traits. In this study, we report the development of linkage map generated with 295 molecular markers. Using this linkage map and accurate phenotyping under drought stress condition we identified seven QTLs associated with drought tolerance. This work provides QTL-based marker-assisted breeding of drought tolerant varieties of horsegram for its genetic improvement.

## Materials and method

### Plant material and experimental design

A RILs population comprised of 162 lines derived from intraspecific crossing of HPKM249 X HPK4 was used in the study. The RIL mapping population was developed through single seed descent method from F_2_ to F_8_ generations. The parents (HPKM249 & HPK4) vary from each other in terms of various drought related traits. HPKM249 cultivar is drought susceptible with less root length and low content of drought related metabolites. HPK4 showed tolerance to drought, long root length and high content of drought related metabolites (Figure 1; Table 1). Parental lines along with 162 RILs were grown in a polyhouse of the Department of Agricultural Biotechnology, CSK Himachal Pradesh Agricultural University Palampur (India) during the summer season of 2017 and 2018. For the phenotypic data for drought related traits, one group of the RILs was planted in a polytubes that were 3 feet long and 4 inches wide. The watering to the plants was stopped at the time of initiation of flowering and the days were counted from day at which watering was stopped till the appearance of drought symptoms. Also these 162 RILs were grown in an augmented block design having two replications in a plot size of 1 m with row-to-row distance of 30 cm between plants and plant-to-plant distance of 10 cm. Four checks (Himganga, HPKC2, VLG1 and HPK4) were grown after every 20 lines in these plots. Five plants were chosen randomly and the data for all the traits were recorded from them and averaged for final readings. Data on days to temporary wilting (DTW), chlorophyll content (CHL), carotenoid content (CAR), Proline content (PRO), MDA content (MDA), relative water content (RWC), membrane stability index (MSI), root length (RTL), root fresh weight (RFW) and root dry weight (RDW) were recorded following standard procedures.

**Table 1.** Morphological polymorphism in parents

| S.No. | Trait | HPK4 | HPKM249 |
| --- | --- | --- | --- |
| 1 | Drought stress | Tolerant | Susceptible |
| 2 | Days to temporary wilting | More | Less |
| 3 | Relative Water Content | High | Low |
| 4 | Total carotenoids | High | Low |
| 5 | Total chlorophyll contents | High | Low |
| 6 | Proline | High | Low |
| 7 | MDA | Low | High |
| 8 | Root length | More | Less |

**Figure 1.**
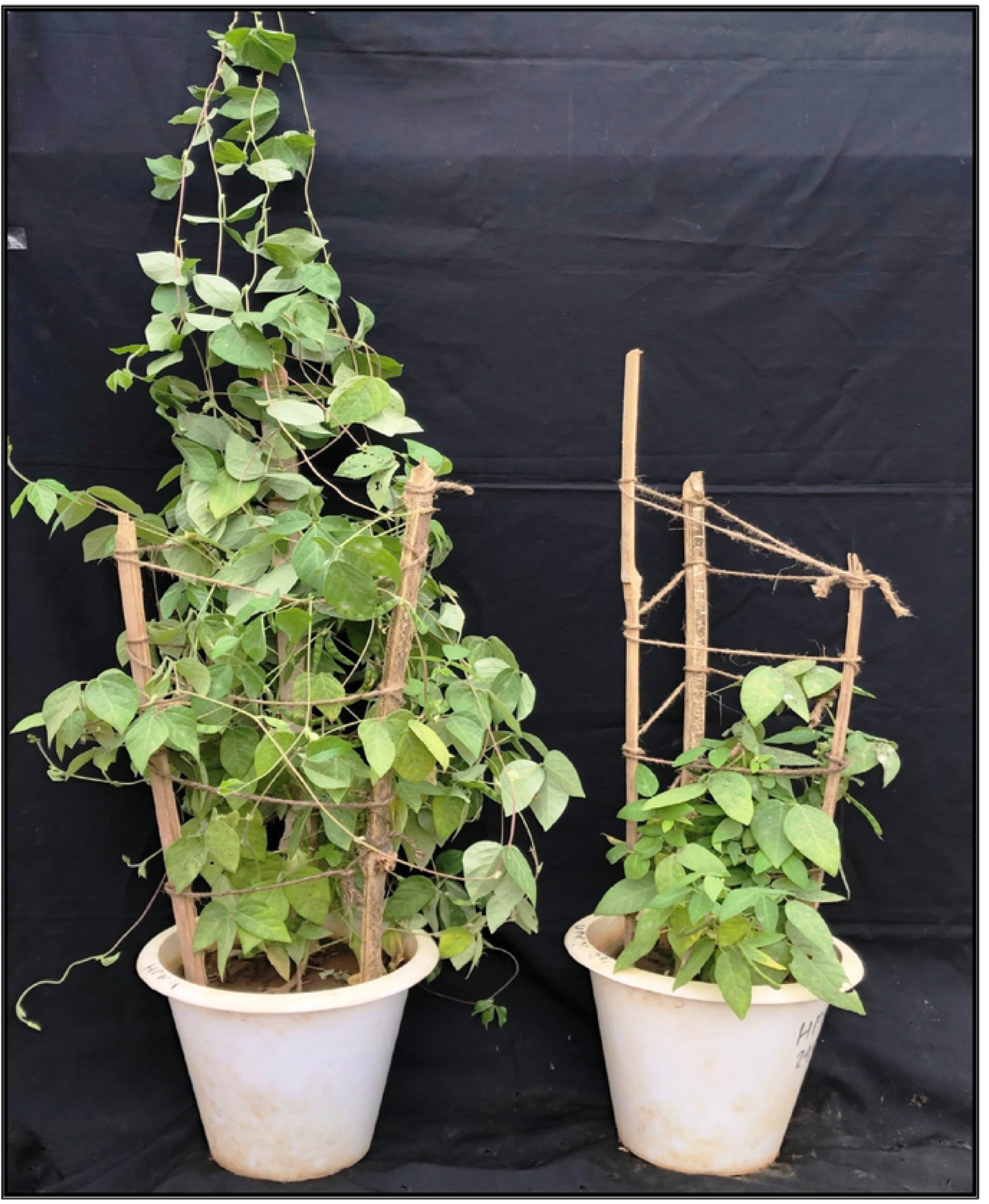
Morphology of the two contrasting parents

### Genotyping

Genomic DNA from young leaf (0.5-1 g) of the parental lines and individuals of F_8_ RILs was isolated using modified CTAB method (Murray and Thompson 1980). DNA concentration was checked by agarose gel electrophoresis. The DNA samples were also quantified on microvolume spectrophotometer (Biospec-nano, Shimadzu Biotech, USA) using Tris EDTA as blank and DNA concentration was recorded in ng/µl.

SSR primers derived from different background were used to detect polymorphism between parental lines. Total of 2011 markers consisting of 63 EST SSRs, 403 horsegram genic SSRs, 387 horsegram genomic SSRs, 24 drought specific SSRs, 300 SSRs from other legumes, 450 RAPD and 384 COS were used for polymorphism survey in parental lines i.e. HPK4 and HPKM249. The polymorphic primer pairs were chosen to genotype all the 162 F_8_ RILs of mapping population. For amplification of genomic DNA, a reaction mixture of 10.0 µl volume was prepared using 4.80µl of sterilized distilled water, 2.0 µl template DNA (13 ng/µl), 0.5 µl of forward and 0.5 µl of reverse primer (5 µM), 0.5 µl MgCl_2_ (25 mM), 1.0 µl 10X PCR buffer (10 mM Tris-Hcl, 50 mM KCl, pH 8.3), 0.5 µl dNTP mix (0.2 mM each of dATP, dGTP, dCTP and dTTP) and 0.2 µl *Taq* polymerase (5U/µl). The amplifications were carried out in Veriti 384^®^ (Applied Biosystems, CA, USA) and 2720 Thermal Cycler (Applied Biosystems, CA, USA). The amplification products were electrophoresed in either 6 per cent PAGE or 3 per cent metaphore agarose gel (Lonza) depending on the resolution pattern, along with size markers. Gels were prepared and run in 1X TAE buffer (3% metaphore agarose gel) or in 1X TBE (6% PAGE) and visualization of fragments was done using Gel-Documentation Unit (ENDURO™ GDS Gel Documentation System, USA) or silver-staining procedure depending upon the requirement. The amplified banding patterns were scored manually as ‘A’ for HPKM249 type banding pattern, ‘B’ for HPK4 and H for heterozygous loci if any. Size of alleles was noted with the help of 100-bp DNA ladder (Fermentas, Lithuania).

### Linkage analysis and map construction

The quantitative trait analysis was done by taking scored data matrix as an input files in JOINMAP® 4.1 program (van Ooijen 2006) for map construction. Minimum independence LOD threshold of 3.0 with a step up of 0.5 till a maximum LOD threshold of 8.0 were used for grouping of markers and to identify linkage groups. Those groups with highest number of mapped markers and maximum linkage at the different LODs were selected. At LOD 4.0, the groups were converted to maps using the regression algorithm. Kosambi’s mapping function was used to calculate the distance (Kosambi 1994).

### Evaluation of drought tolerance

Phenotyping and evaluation for ten drought related traits by using the mapping population of 162 RILS for two consecutive summers (2017 and 2018) at Palampur (H.P.), India (32.1167^0^ N, 76.5333^0^ E; 1220 m.a.s.l.) was done. For phenotypic data, we planted RILs in a polytube that was 3 feet long and 4 inches wide. At the time of flower initiation, the watering was stopped and the days were counted to the appearance of drought symptoms which is recorded as days to temporary wilting (DTW). The chlorophyll content (CHL) was estimated using Yoshida et al. (1976), the amount of carotenoids (CAR) was determined according to Lichtenthaler and Wellburn (1983), the free proline content (PRO) was estimated by the method of Bates et al. (1973), the degree of lipid peroxidation was measured in terms of MDA content as described by Heath and Packer (1968) from both well watered and drought stressed RIL population along with parents. Relative water content (RWC) was measured on the second or third upper fully expanded leaf for both well-watered and stressed plants. It was calculated using the formula of Shrestha et al. 2006 as RWC (%) = [(FW − DW)/ (TW − DW)] × 100, where FW is the leaf fresh weight, DW is the leaf dry weight after oven drying (80 °C) for 48 h; and TW is the leaf turgid weight (obtained after rehydration of the leaf samples in distilled water for 4 h). MSI was estimated according to Sairam (1994). In this, two sets of leaf tissues (0.1 g) were placed in 10 ml of double distilled water. One set was kept at 25°C for 24 h, kept on shaking, initial conductivity (Ci) of the bathing solution was measured with the conductivity meter. Second set of tissue was autoclaved at 121°C for 30 min and cooled down to 25°C before final conductivity (Cmax) was measured as MSI (%) = 1 – (electric conductivity before incubation/ electric conductivity after incubation). Root length (RL, cm), root fresh weight (RF, g) and root dry weight (RD, g) was also calculated at the time of harvesting by uprooting and removing plant and soil carefully. All recommended agronomic practices were followed during the cropping season.

### Phenotypic data analysis and QTL mapping

Quantitative trait loci analysis was done on 162 RILs using phenotypic data of different drought related traits evaluated in the study. Distribution of each trait was inspected using skewness and kurtosis statistics using Past 3.25 software. Phenotypic differences between parents were analyzed using ANOVA. Pearson correlation coefficients and frequency distribution among different traits were calculated using the same software. QTLs were identified using Windows QTL Cartographer V2.5 software (Wang et al. 2005) by composite interval mapping (CIM) method (Zeng 1993; 1994). The Zmapqtl standard model 6 was used with a 10 cM of window size and walk speed of 2 cM. Cofactors were obtained using the forward regression algorithm. A LOD score threshold of a QTL at a significance level of P=0.05 was estimated at 1000-permutation of shuffling the genotypes and means of phenotypic values for a trait (Doerge and Churchill 1996). An LOD threshold score of ≥2.5 at 1000 permutations was significantly considered (5% level of significance) to identify and to map the QTLs on the horsegram LGs. The 95% confidence intervals of the QTL locations were determined by one LOD intervals surrounding the QTL peak (Mangin et al. 1994). The estimated additive effect and the percentage of phenotypic variation explained by each putative QTL were obtained using the software with the CIM model by the Zmapqtl procedure. The R^2^ value from this analysis was accepted as the percent phenotypic variance explained by the locus. The figure of the qtl map was made with MapChart 2.32 software (Voorrips 2002).

## Result

### Parental screening and genotyping of mapping population

The polymorphism survey was done on parental lines using 1177 SSR primer pairs [63 (Horsegram EST SSRs) + 403 (Horsegram genic SSRs) + 387 (Horsegram genomic SSRs) + 24 (drought specific SSRs) + 300 (SSRs from other legumes viz. red clover and *Medicago*)] of which 430 were found to be polymorphic. Along with this, 450 RAPD primers were screened out of which 55 were found to be polymorphic and also 384 COS markers of *Medicago truncatula* were screened out of which 8 were found to be polymorphic. Total of 2011 primers were screened, out of which 493 were found to be polymorphic and these were then used for genotyping of the 162 individuals. The size of amplified bands produced by 493 polymorphic primers ranged between 100 and 250 bp. The genotyping data thus obtained was scored manually and used as the input file for the construction of horsegram linkage map. The total number of markers screened and the number of polymorphic markers obtained has been listed in Table 2.

**Table 2.**
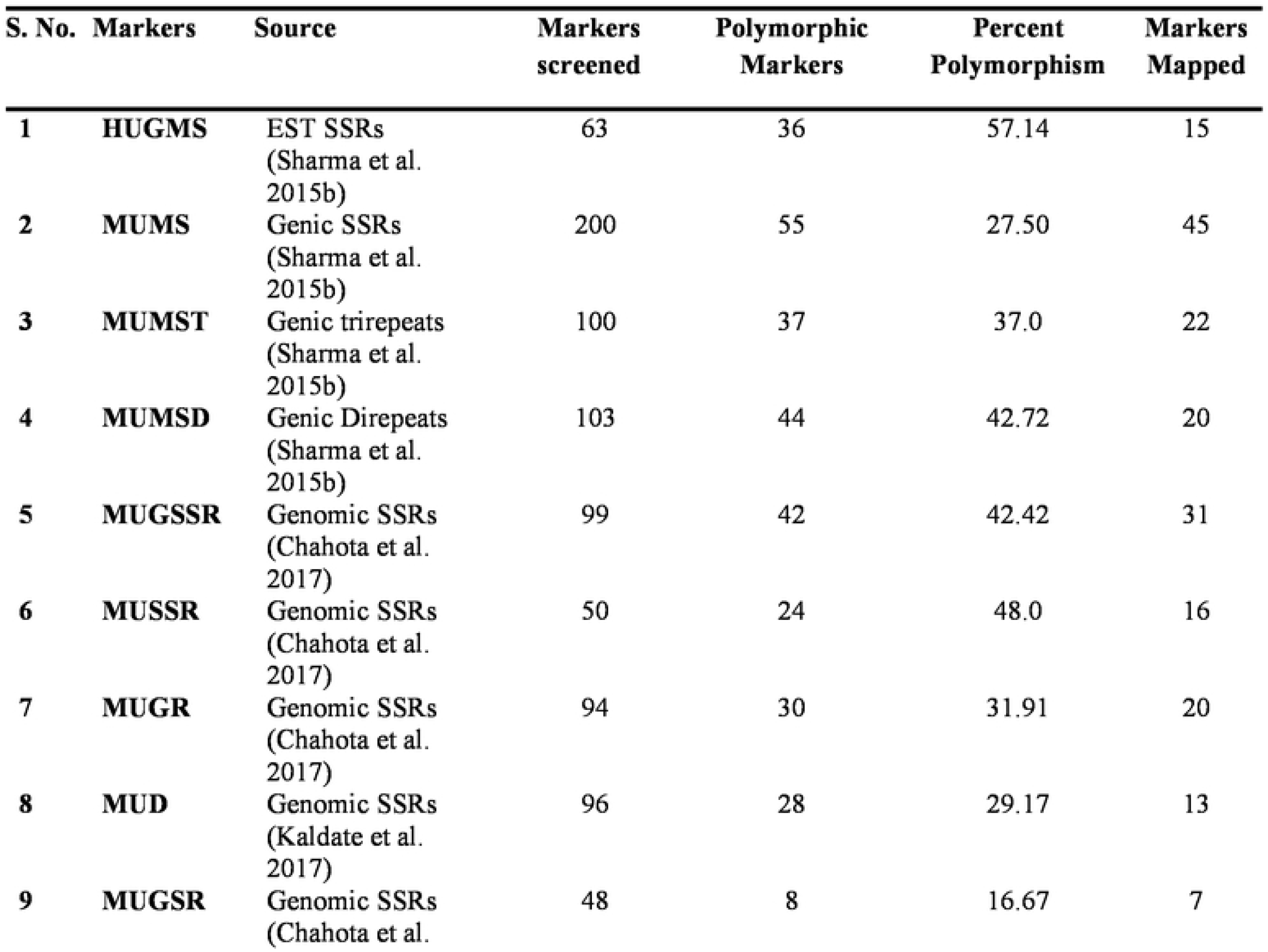
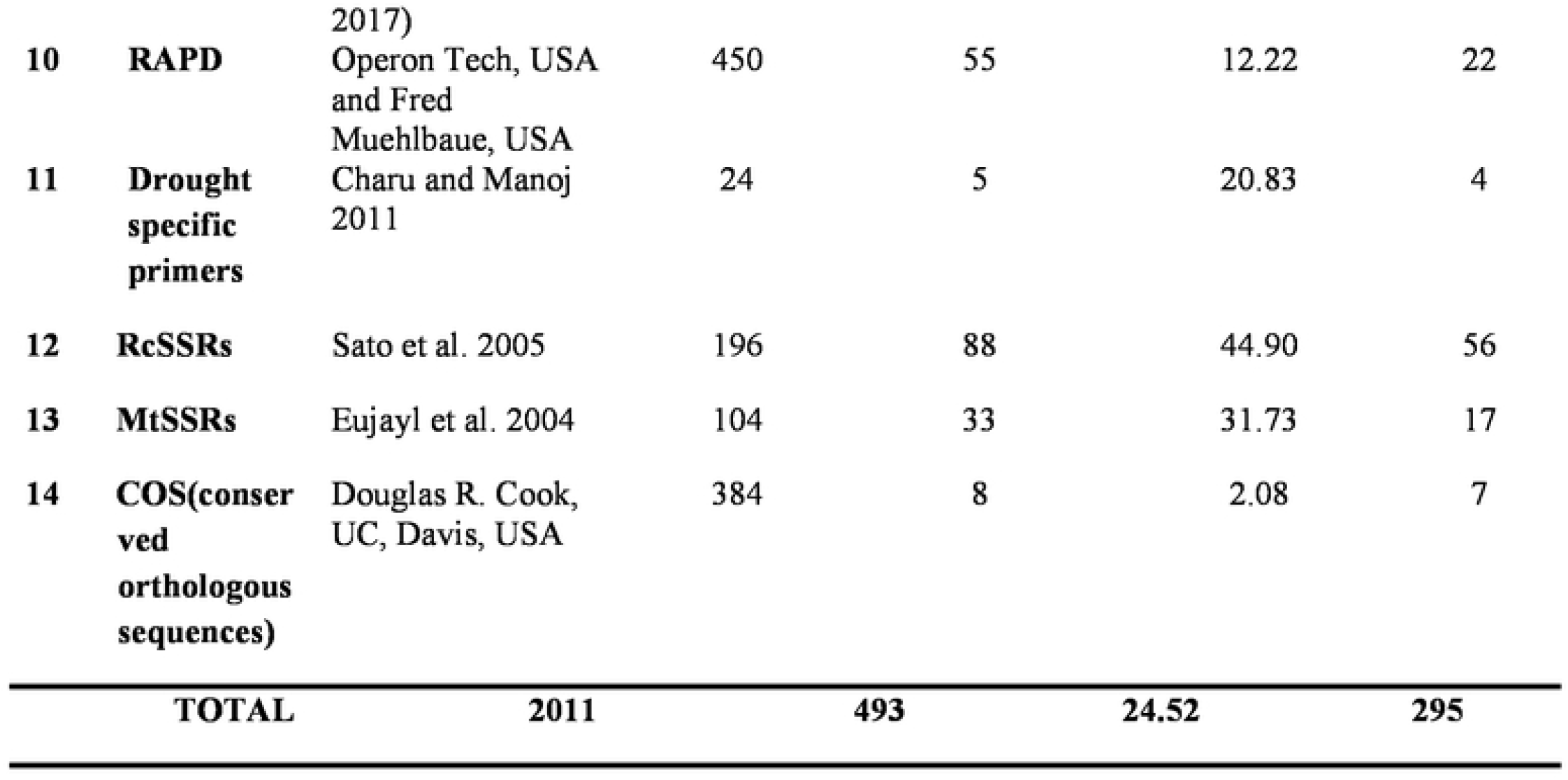
Markers used for construction of intra-specific linkage map of horsegram

### Construction of genetic linkage map

Of the 493 polymorphic markers, 295 (59.84%) were mapped into 10 LGs at LOD 3.5 using JoinMap software, version 4.0. The map spanned 1541.7 cM (Kosambi cM) length and an average marker interval size of 5.20 cM (Table 3). Of these 295 mapped markers include, 15 EST SSRs, 87 genomic SSRs, 87 genic SSRs, 22 RAPDs, 73 SSRs from other species, 4 drought specific markers and 7 COS SSRs. Each of the ten linkage groups differed from one another in terms of their length and total number of marker mapped. Of the total 295 mapped markers, LG1 harboured 89 markers followed by LG2 which contained 58 markers. 35 markers were mapped on LG3, 29 markers were mapped on LG4, 19 were mapped on LG5 and LG7 whereas 18 markers were mapped on LG6. LG 8 and LG 9 contained least number of markers with 7 and 6 markers respectively and linkage group 10 harboured 15 markers. Though the LG7 is having the maximum size of 238.5 cM distance but due to very less number of markers (19) present on it with average marker density of 12.5 however is of less importance while mapping of different QTLs in comparison to linkage groups LG1 and LG2 having marker density of 2.0 and 2.7 respectively even having smaller linkage size (182.9 cM and 159 cM). Segregation distortion for all the 493 polymorphic markers was determined and of these 295 (59.83%) followed the expected segregation ratio, whereas 198 markers (40.16%) were found to show deviation. These distorted markers were excluded from the final analysis.

**Table 3.** Distribution of 295 markers on ten linkage groups of an intra-specific linkage map of horsegram.

| **LGs** | **Markers Mapped** | **Map length (cM)** | **Average marker density (cM)** |
| --- | --- | --- | --- |
| **LG1** | 89 | 182.9 | 2.0 |
| **LG2** | 58 | 159.0 | 2.7 |
| **LG3** | 35 | 129.0 | 3.7 |
| **LG4** | 29 | 188.0 | 6.5 |
| **LG5** | 19 | 192.5 | 10.1 |
| **LG6** | 18 | 165.6 | 9.2 |
| **LG7** | 19 | 238.5 | 12.5 |
| **LG8** | 7 | 71.6 | 10.2 |
| **LG9** | 6 | 46.4 | 7.7 |
| **LG10** | 15 | 168.2 | 11.2 |
| **Total** | **295** | **1541.7** | **5.2** |

The maximum distance between markers was 61.2 cM on LG7 and the minimum distance was 0.003 cM on LG2. The number of markers present in different linkage groups was unequal. Four large groups having 12-19 markers within a length of 10 cM and five groups having 28-31 markers within a length of 30 cM was found. The length of the linkage groups did not reflect the number of markers linked on it as the distance between markers varied across different linkage groups. For example, LG1 carrying 89 markers having a length of 182.9 cM and an average marker distance of 2.0 cM, whereas LG4 having a length of 188.0 cM carrying 29 markers and average marker distance of 6.5 cM and LG5 having a length of 192.5 cM covered by 19 markers and average spacing of 10.0 cM.

### Phenotypic Evaluation of Parents and the Mapping Population for Drought Tolerance-Related Traits

In the present study ten drought related traits were analysed namely days to temporary wilting (DTW), chlorophyll content (CHL), carotenoid content (CAR), Proline content (PRO), MDA content (MDA), relative water content (RWC), membrane stability index (MSI), root length (RTL), root fresh weight (RFW) and root dry weight (RDW) were analyzed. The two parents, HPKM249 and HPK4 used to develop mapping population showed significant differences in the drought related traits. HPK4 has higher value for days to temporary wilting (DTW), chlorophyll content (CHL), carotenoid content (CAR), Proline content (PRO), membrane stability index (MSI), root length (RTL), root fresh weight (RFW) and root dry weight (RDW) and thus is drought tolerant whereas HPKM249 has higher value for MDA content (MDA). All the traits showed continuous distribution on histogram and exhibited transgressive segregation which is an important characteristic of quantitative traits (Figure 2). Descriptive statistics for drought related traits in the RIL population is presented in Table 4. Results showed that all the lines response differently to water stress for all traits studied. All the traits were significantly affected by drought stress. Phenotyping for these traits showed significant genetic variability within RILs population under normal and water stressed condition. Correlations among traits calculated within RILs revealed that root traits were found positively correlated to each other and were statistically significant (P<0.05). Other drought traits were also positively correlated to each other and were statistically significant (P<0.05) under different environment (Figure 3).

**Table 4.** Mean performance of parents and RILs at Palampur 2017 for biochemical and root traits in RIL (HPKM249 × HPK4) population

| Traits | Environment | HPKM249 | HPK4 | Range (RIL) | Mean | SD |
| --- | --- | --- | --- | --- | --- | --- |
| **Biochemical** |  |  |  |  |  |  |
| Carotenoids | Control | 0.81 | 0.78 | 0.11-0.99 | 0.56 | 0.17 |
|  | Stress | 0.58 | 0.65 | 0.06-0.99 | 0.48 | 0.21 |
| Chlorophyll | Control | 11.46 | 12.97 | 4.07-24.64 | 13.36 | 5.09 |
|  | Stress | 7.91 | 10.59 | 1.69-22.58 | 11.00 | 5.01 |
| Malondialdehyde | Control | 16.84 | 13.47 | 10.24-19.78 | 15.22 | 2.27 |
|  | Stress | 29.65 | 20.83 | 17.15-32.59 | 25.23 | 3.53 |
| Proline | Control | 0.21 | 0.46 | 0.07-0.97 | 0.45 | 0.22 |
|  | Stress | 2.03 | 3.43 | 1.14-3.65 | 2.20 | 0.59 |
| Membrane stability index | Control | 0.94 | 1.00 | 0.80-1.00 | 0.94 | 0.05 |
|  | Stress | 0.86 | 0.93 | 0.69-1.00 | 0.85 | 0.07 |
| Relative water content | Control | 87.34 | 96.36 | 83.39-98.69 | 89.93 | 3.56 |
|  | Stress | 74.44 | 86.58 | 65.34-89.58 | 77.68 | 5.00 |
| **Root traits** |  |  |  |  |  |  |
| Root length | - | 51.00 | 63.00 | 40.00-89.00 | 60.75 | 8.26 |
| Root fresh weight | - | 0.42 | 1.34 | 0.24-6.26 | 2.66 | 1.47 |
| Root dry weight | - | 0.08 | 0.97 | 0.04-5.19 | 1.96 | 1.27 |

**Figure 2.**
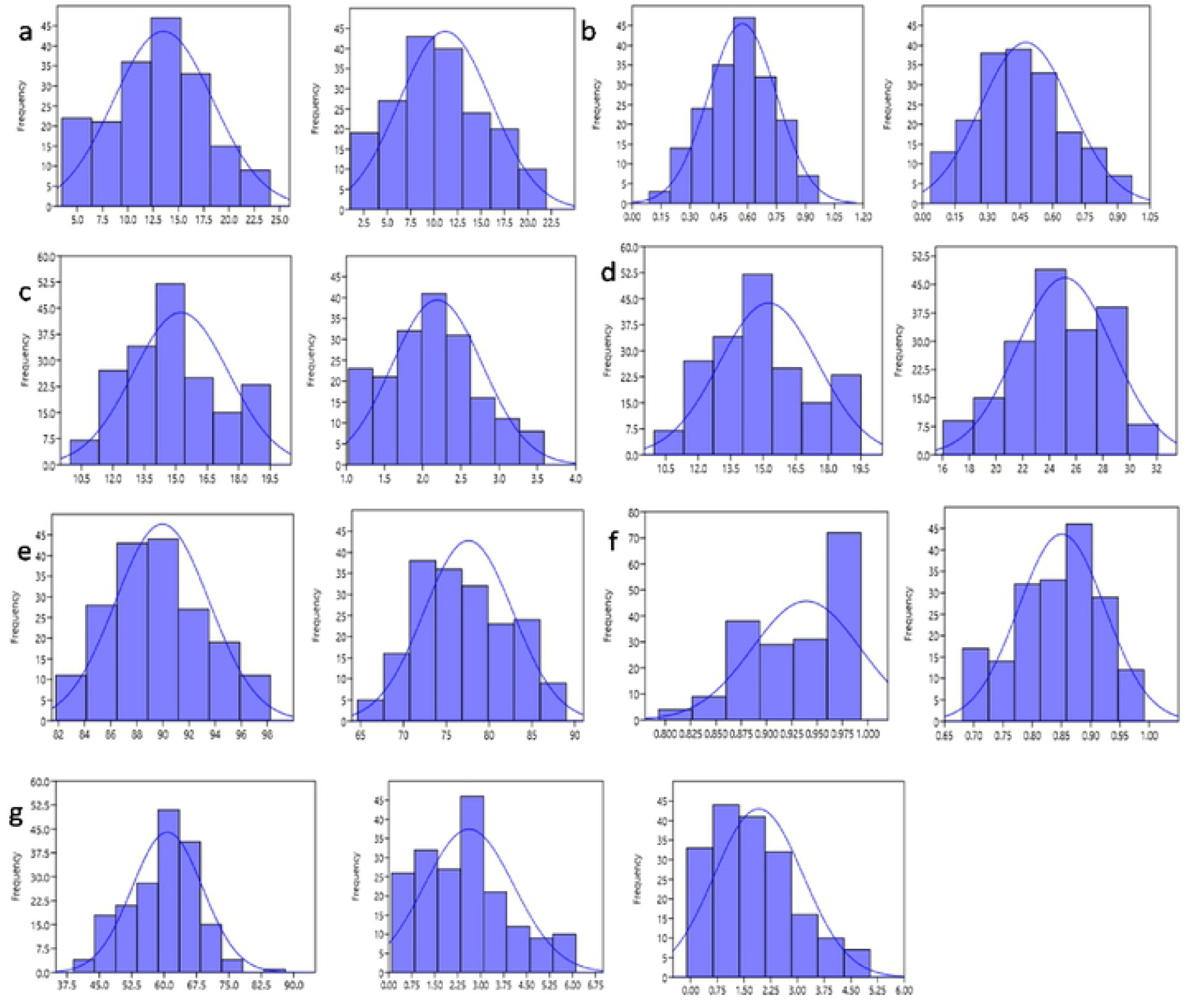
Frequency distribution curve of (a) Chlorophyll content in control and stress condition (b) Carotenoid content in control and stress condition (c) Proline content in control and stress condition (d) MDA content in control and stress condition (e) RWC in control and stress condition (f) MSI in control and stress condition (g) Root length, Root fresh weight and Root dry weight

**Figure 3.**
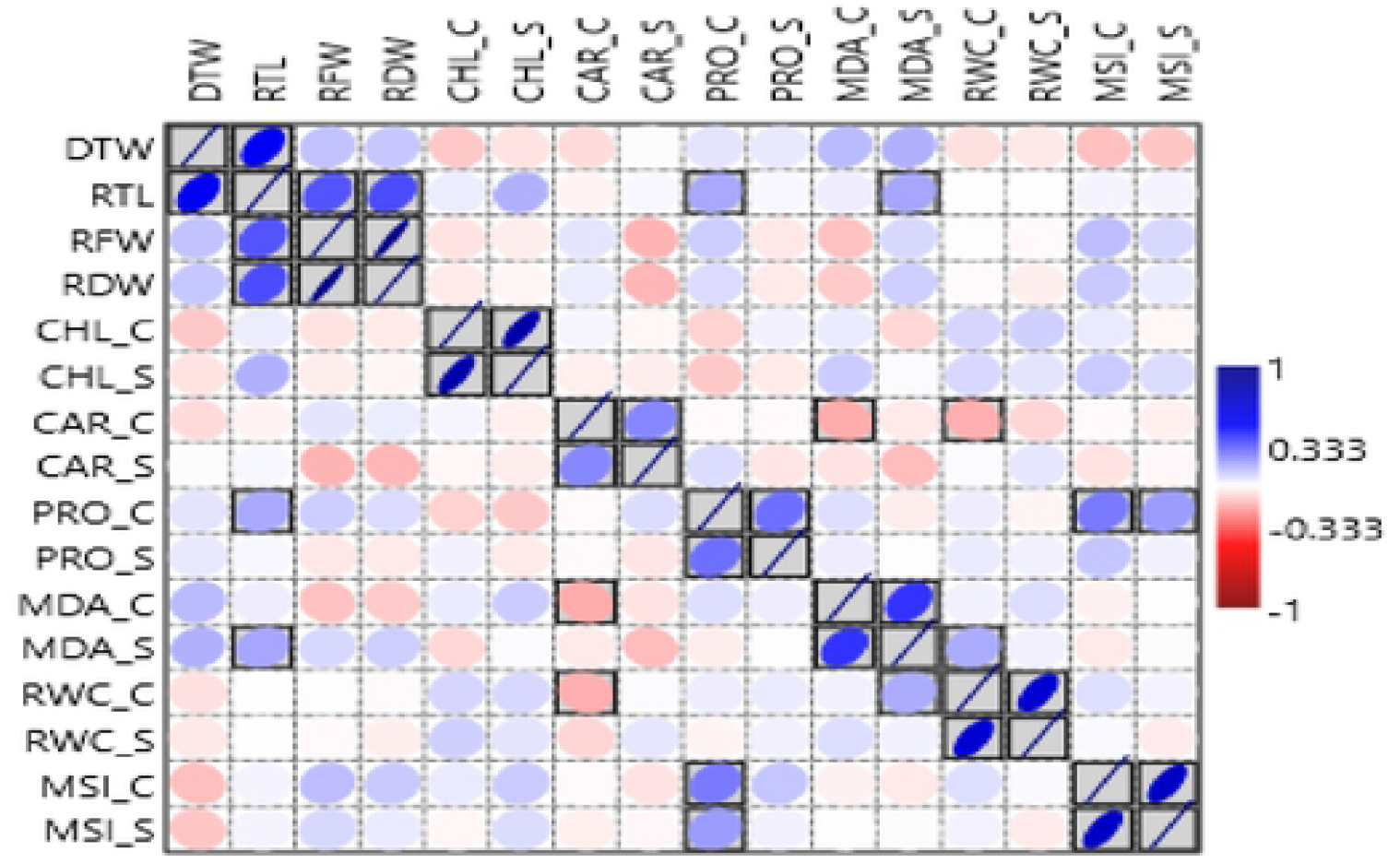
Pearson’s correlation matrix among different traits analyzed in the HPM249 × HPK4RILs

### QTL Mapping

QTLs for the different drought related traits were identified by QTL Cartographer V2.5 using phenotypic data of RILs under normal and water stressed condition. Total of seven QTLs were identified for ten different drought related traits namely one for chlorophyll content under drought stress, one for malondialdehyde content for control and one for proline content under drought stress, one for root dry weight, one for root fresh weight and two for root length using composite interval mapping (Table 5).

**Table 5.**
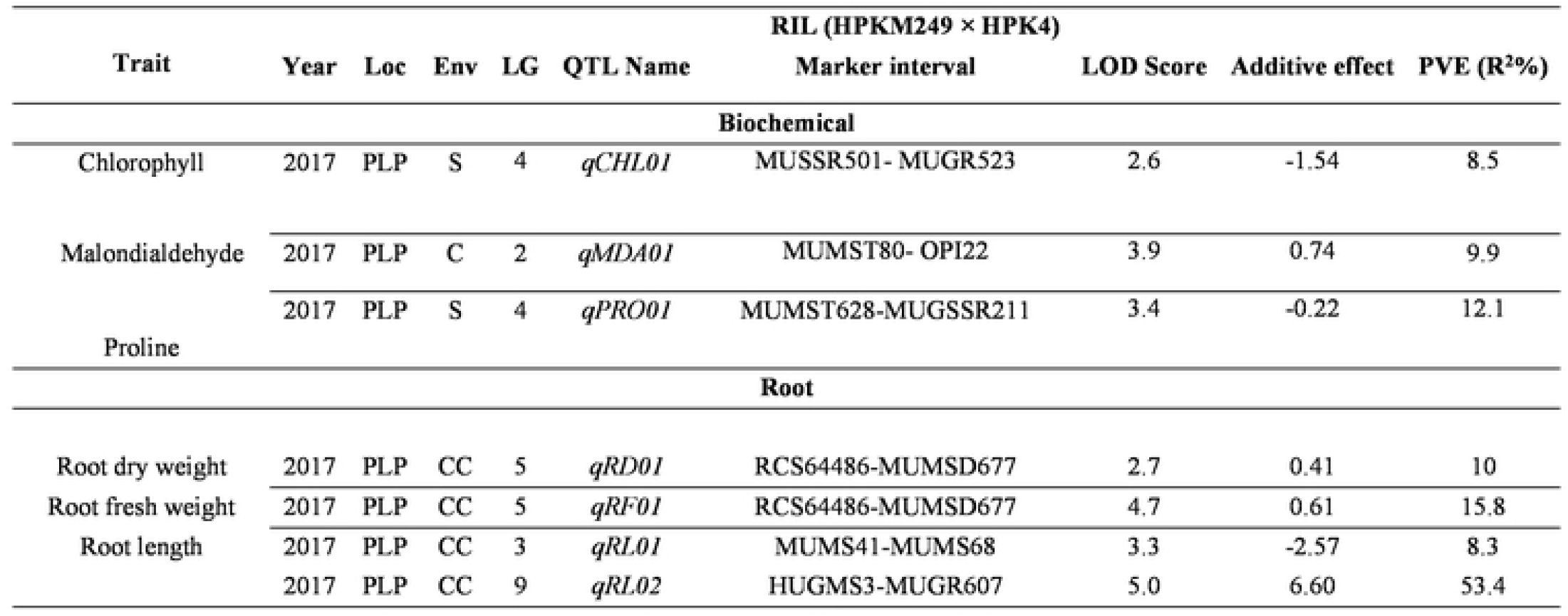
QTLs for various drought related traits identified using QTL Cartographer

One drought specific QTL for chlorophyll content was present on LG4 with phenotypic variation explained of 8.5 per cent at LOD value of 2.6 flanked by MUSSR 501-MUGR 523 marker interval. Similarly, one drought specific QTL was also detected for proline content at LOD of 3.4 with 12.1 per cent of phenotypic variation explained. This QTL was also present on LG4 flanked by MUMST628-MUGSSR211 marker interval. One QTL for malondialdehyde content was present on LG3 with 9.9 per cent of phenotypic variation explained at LOD 3.4 flanked by MUMST80-OPI22 marker interval. Additive effect demonstrated that HPK4 contributed alleles for chlorophyll, proline and HPKM249 contributed alleles for malondialdehyde content. QTL for root dry weight was present on LG5 with 10 per cent of phenotypic variation explained at LOD value of 2.7, flanked by RCS64486-MUMSD677 marker interval while QTL for root fresh weight was also present on LG5 with 15.8 per cent of phenotypic variation explained at the LOD value of 4.7, also flanked by RCS64486-MUMSD677 marker interval. Both these QTLs were contributed by the alleles from the parent HPKM249 which resulted in increased root dry weight and root fresh weight. Two QTLs were detected for root length at LOD of 3.3 and 5.0 with 8.3 and 53.4 per cent of phenotypic variation. These QTLs were present on LG3 and LG9 flanked by MUMS41-MUMS68 and HUGMS3-MUGR607 marker interval, respectively. Further, additive effect demonstrated that allelic contribution is by both the parents resulted in increased root length (Figure 4).

**Figure 4.**
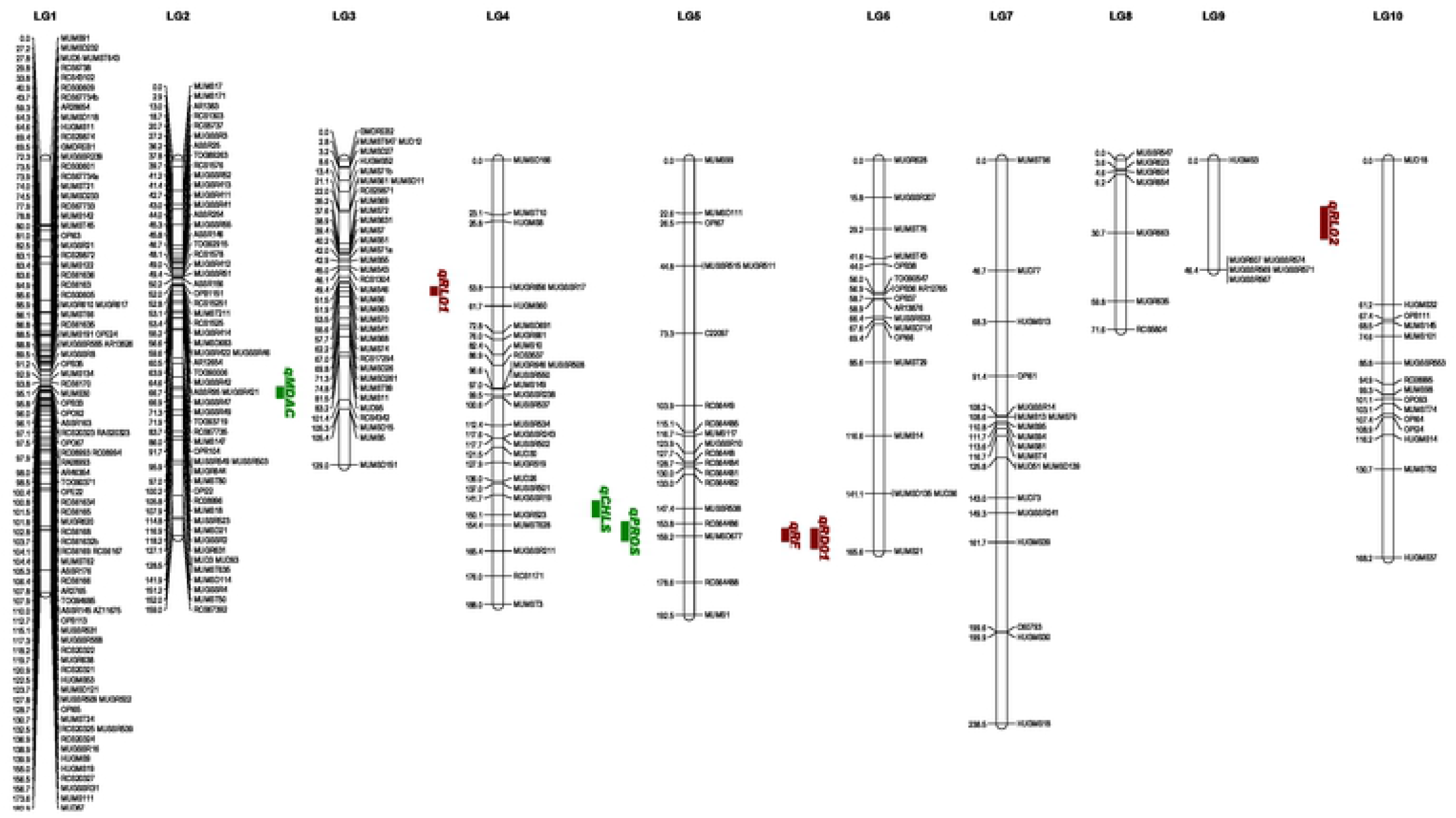
Likelihood intervals for quantitative trait loci (QTLs) associated with drought related traits in recombinant inbred lines (RILs) mapping population

**Figure 5.**
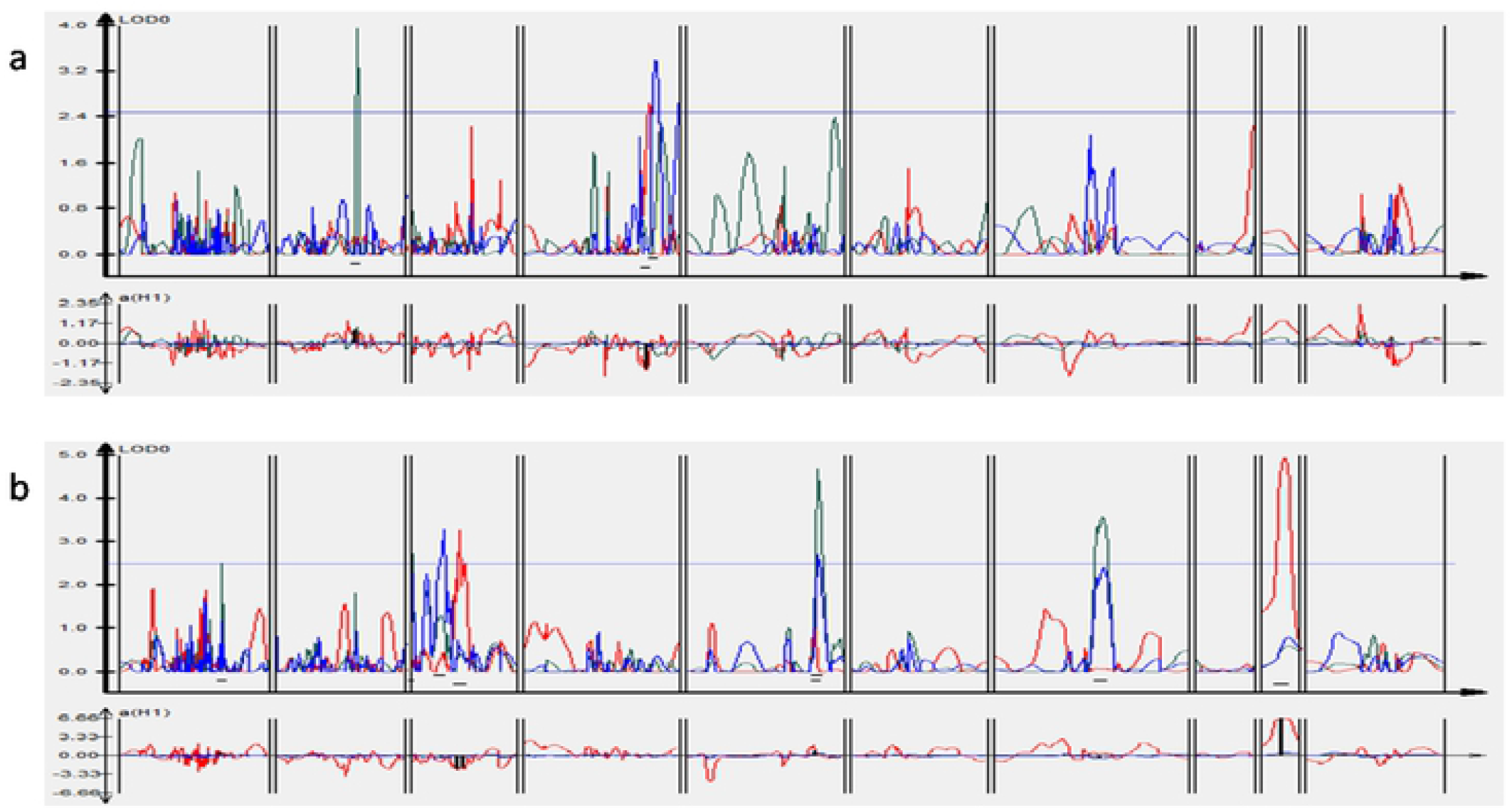
(a) Position of QTLs for (a) biochemical traits (b) root traits on 10 linkage groups of horsegram

## Discussion

Horsegram is one of the important crop in terms of its medicinal, neutracutical and drought tolerant properties but ignored by the scientific community in terms of its genetic improvement. It is mainly grown by local people as ethnical crop in some areas of developing countries (Chahota et al. 2013; Fuller and Murphy 2018). It is widely grown as food for providing balanced nutrition to resource poor people and as feed for animals in dry areas. Also medicinal, neutraceutical and environmental advantages make horsegram an important food source of future.

For genetic improvement of any crop, construction of linkage maps is prerequisite which will leads to mapping of important agronomic traits such as yield and resistance to various biotic or abiotic stresses. Availability of large number of molecular markers is essential for the construction of linkage map and other plant molecular analysis. Knowledge of plant genomic research has increased manifolds due to the development of molecular markers. These DNA molecular markers aids in increased reliability and resolution of mapping. However the information of molecular markers in horsegram is very limited but few attempts have been made in last decade (Sharma et al. 2015a, b; Chahota et al. 2017; Kaldate et al. 2017). Therefore for the construction of a high density linkage map in horsegram, more molecular markers were developed and used in the study. These SSRs were developed from EST sequences, data from transcriptome sequences (Bhardwaj et al. 2013) and data from Illumina DNA sequencing (Illumina Inc., San Diego, CA, USA)000000000. Also markers from other legumes were used in the study.

To evaluate the efficiency of the developed primers for their amplification, all the primers developed were screened on the parental lines (HPKM249 and HPK4) for polymorphism analysis.

In the study total of 2011 markers consisting of 63 EST SSRs, 403 genic SSRs, 387 genomic SSRs, 24 drought specific SSRs, 300 SSRs from other legumes, 450 RAPD and 384 COS were used for polymorphism survey on the parental lines of the RIL mapping population i.e. HPK4 and HPKM249. Of these, 493 (24.51) were found to be polymorphic which comparable to other legumes. The polymorphism shown by the mapping population relies on many aspects such as type of marker and population.

A total of 294 markers were mapped on ten linkage group covering a distance of 1541.7 cM (Kosambi cM) with an average marker density of 5.20 cM. Since SSRs are present randomly on the genome and due to biasness towards selection of genomic region for the identification of SSRs each linkage group exhibited different length and distribution of marker. Therefore LG1 and LG3 have densely distribution of markers, while LG7 and LG6 have only few markers on them. The markers were ordered on a particular linkage group on the basis of minimum level of confidence at LOD 3.0. After the construction of Linkage map, it can be utilized for the mapping of important traits such as yield, biotic and abiotic stress tolerance, etc.

Out of numerous stresses, plant suffer huge lose in its production due to drought (Malhotra et al. 2004; Singh et al. 1994). To increase the production and ultimately the profit from legumes drought tolerance is an important traits (Idrissi et al. 2015). Therefore to identify and locate the genes in the genome which can directly influence the drought tolerance is an important criteria to ease the selection of lines in breeding programme through MAS. Since drought tolerance is a complex trait and is controlled by many genes directly and indirectly, accurate phenotyping and analysis of several biochemical and physiological traits which affects it need to be investigated on the mapping population. Also study on root traits can provide better insight as roots are the first organ which are exposed to drought stress in plants. In the present study, the F8 RIL mapping population was evaluated for different nine drought related traits for two consecutive years in different environments. Using composite interval mapping, a total of seven QTLs were detected viz. one for each chlorophyll content under drought stress, malondialdehyde content under contol and proline content under drought stress, root dry weight, root fresh weight and two QTLs for root length. LG2 contained 1 QTL for MDA under control condition, LG3 contained 1 QTL for root length, LG4 had a total of 2 QTLs one each for Chlorophyll and Proline both under stress condition, LG5 contained a total of 2 QTLs one each for root dry weight and root fresh weight and LG9 contained 1 QTLs for root length. Studies in other crops also confirm the uneven localization of QTLs across the linkage group (Zhang et al. 2015, Pottorff et al. 2014, Leite et al. 2011 and Fratini et al. 2007). Addition of more molecular markers such as SNPs to the linkage map can further provide better localization and estimation of QTLs, which can be used for the MAS and genetic improvement of the horsegram.

## Contribution statement

RKC planned study and finalized manuscript, MK performed the experiment, recorded data and performed molecular data analysis.

